# FABEL: Forecasting Animal Behavioral Events with Deep Learning-Based Computer Vision

**DOI:** 10.1101/2024.03.15.584610

**Authors:** Adam Catto, Richard O’Connor, Kevin M. Braunscheidel, Paul J. Kenny, Li Shen

## Abstract

Behavioral neuroscience aims to provide a connection between neural phenomena and emergent organism-level behaviors. This requires perturbing the nervous system and observing behavioral outcomes, and comparing observed post-perturbation behavior with predicted counterfactual behavior and therefore accurate behavioral forecasts. In this study we present FABEL, a deep learning method for forecasting future animal behaviors and locomotion trajectories from historical locomotion alone. We train an offline pose estimation network to predict animal body-part locations in behavioral video; then sequences of pose vectors are input to deep learning time-series forecasting models. Specifically, we train an LSTM network that predicts a future food interaction event in a specified time window, and a Temporal Fusion Transformer that predicts future trajectories of animal body-parts, which are then converted into probabilistic label forecasts. Importantly, accurate prediction of food interaction provides a basis for neurobehavioral intervention in the context of compulsive eating. We show promising results on forecasting tasks between 100 milliseconds and 5 seconds timescales. Because the model takes only behavioral video as input, it can be adapted to any behavioral task and does not require specific physiological readouts. Simultaneously, these deep learning models may serve as extensible modules that can accommodate diverse signals, such as in-vivo fluorescence imaging and electrophysiology, which may improve behavior forecasts and elucidate invervention targets for desired behavioral change.

## 1 Introduction

A key aim of behavioral neuroscience is to understand the individual and collective roles of brain regions, neural circuits, and specific cell types implicated in neurobehavioral disorders, in order to discover novel interventions. The goal of applying these interventions is to effect measurable change in relevant behavioral endpoints; thus a proper quantification of such behaviors is critical to empircally determine whether interventions have had significant effects. Laboratory animal models of neurobehavioral disorders provide an effective window for study, with the ability to (i) interrogate their underlying biological processes in real-time via calcium imaging, fiber photometry, electrophysiology, and behavioral video recording; (ii) selectively control biological state via transgenic Cre drivers, CRISPR-mediated gene knockouts, and behavioral training; (iii) apply real-time interventions such as foot shocks, audible tones, and optogenetic or electrophysiologic stimulation of individual cell types, circuits, or brain regions. The typical way to assess the effect of interventions on behavioral endpoints is to compare the distributions of occurrences of the behavioral endpoints across condition and control groups. While this paradigm provides insight into the effects of interventions on global tendency to engage in behavior, it is agnostic to individualized context and therefore does not allow for (1) personalized prediction of behavior change, and (2) prediction of real-time trial-by-trial event occurrences in response to interventions. Assessing the effects of interventions on real-time behavior requires accurate counterfactual behavior, i.e. forecasts of future behaviors in the absence of interventions. Behavior forecasting models could be incorporated with closed-loop optogenetic and electrophysiologic stimulation systems to determine optimal intervention schedules to have a desired effect on behavioral endpoints while minimizing side effects.

One modality that holds rich information for behavior forecasting is in vivo neural activity. There is extensive literature documenting the correlation between neural dynamics and behaviors of interest. One exemplary research area is motor planning, in which cortical electrical activity is correlated with motor behaviors such as limb reaching [1, 2, 3, 4, 5], which is important for control of neural motor prosthetics [6, 7]. Another line of research concerns the prediction of future decision-making [8, 9] and goal-oriented behavior [10, 11, 12, 13]. Much work has been done on neural and physiological decoding for future behavior prediction as a component of closed-loop neuromodulation in Parkinson’s Disease, specifically for the prediction of tremor and freezing of gait [14, 15, 16, 17]. These studies help advance our understanding of patterns of neural activity that generate action sequences and drive decisions, thus helping to predict future behaviors; however, they typically focus on simple neural activity patterns that precede the onset of a behavior, such as thresholding and population activity that can be linearly decoded. The behaviors tend to be simple well-defined movements or simple decision-making tasks, in very controlled settings in which motion and actions are highly restricted. Aside from neural decoding for motor prosthetics and neuromodulation, the approaches in these behavior prediction studies have not been implemented in real-time behavior prediction systems, namely for goal-oriented and otherwise complex decision-making tasks. Furthermore, there are many potential readout modalities that can be recorded in a behavioral experimental setting; aside from neural (e.g. calcium imaging, in vivo electrophysiology, fiber photometry) and physiological (e.g. wearable metabolic sensors [18], wearable ECG monitor [19]) recording, **behavioral video** presents a rich stream of data that can reflect important information about an animal’s biological and cognitive state and dynamics. Behavioral video is an especially desirable modality due to its relatively low cost in terms of money and setup time, requiring only a video camera. Many sophisticated tools have been developed to compress, analyze, and interpret animal behavioral video, from animal pose estimation [20, 21, 22] to behavioral understanding [23, 24, 25, 26, 27, 28]. There is a large ecosystem of open-source tools for behavioral video analysis [29, 30] and behavioral experiment automation [31, 32]. Such tools allow researchers to quantify and interpret complex behaviors from large datasets in an automatic or semi-automatic way, yielding new insights and experimental capabilities at scales unimaginable without machine learning and automation. Despite the widespread use of these tools in the behavioral neuroscience community, there is no existing research attempting to forecast future behaviors from behavioral video data. There is a large machine learning literature dedicated to time series forecasting in the context of human activity, from forecasting of future human poses [33, 34, 35] to prediction of specific activities [36, 37, 38, 39, 40, 41, 42, 43, 44, 45], as well as prediction of future semantic information in video [46]. Given that locomotion patterns are implicated in – and defining criteria of – neurobehavioral disorders [47, 48], it is useful to predict future behaviors from behavioral video, both in order to understand the antecedent patterns that yield adverse (or favorable) behaviors and to automatically intervene in the experiment in real-time.

We present a proof-of-concept demonstrating the predictability of behavioral events from locomotion patterns in behavioral video alone. Herein we develop a deep learning-based computer vision approach for forecasting user-defined behavioral events from a behavioral video stream at varying timescales in real-time. Specifically, we forecast a mouse’s interactions with food (e.g. sniffing, eating) in an open field with lag-times between 100 milliseconds and 5 seconds. Accurate forecasts of an animal’s baseline food-seeking behavior provides a counterfactual basis for studying the effects of interventions in the context of obesity, and the approach can translate to other goal-oriented behaviors such as drug-seeking. Our approach leverages fast animal pose estimation as a preprocessing step to reduce the dimensionality of the behavioral videos. A deep neural network sequence model is trained to forecast the likelihood of behavioral event occurrence from sequences of poses. Importantly, this system can be used as a component in closed-loop behavioral experiment platforms, both reducing manual labor and probing understanding of how to prevent the onset of behavioral disorder phenotypes.

## 2 Methods

### 2.1 Dataset Construction

Two-dimensional animal behavior videos were recorded at a resolution of 320 *×* 240 pixels, at a sampling rate of 30 FPS. To compress the dimensionality of the videos, we used pose estimates of four body parts: snout, left-ear, right-ear, and mid-back. We annotated between several dozen and several hundred frames for each video, and trained a separate pose estimation network using the SLEAP software package [22] for each video. The outputs of the pose estimation networks are sequences of (*x, y*) coordinates of each body part. In our experiments, the videos are between 10 *−* 30 minutes and capture mouse feeding behavior in an operant chamber. A bowl of chow is set near the middle of the frame; for each video, we note the (*x, y*) pixel coordinates of the center of the bowl, as well as the radius of the bowl. The body part coordinates in each video are registered to the bowl’s center by subtracting the bowl center coordinates *c* from each body part’s (*x, y*) coordinates, then dividing by the video’s resolution dimensions. Mathematically, we define this coordinate transformation as:

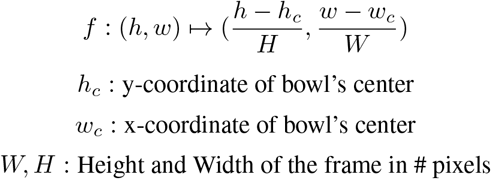

This way all videos are embedded into a common coordinate system with the bowl’s center as the origin, and normalized to the video’s resolution. In this study, each timestamp is provided a label as either interacting with or not interacting with the food, automatically generated by whether or not the snout is positioned within the bowl’s radius. The result is a sequence of pose vectors aligned to the center of the food bowl, with corresponding food-interaction labels at each timestamp. These sequences constitute the input domain to the forecasting network.

### 2.2 Neural Network Architecture

The forecasting network is a deep LSTM recurrent neural network architecture, which maps a “history” of pose vectors to a future behavioral label. Given an *n*-dimensional pose vector, a history length *L*, a batch size *B*, and a future target label ℒ to predict, the forecasting LSTM network is a function *f* :< *L, n* > → ℒ. The actual target label is task-dependent, and in general is based on the historical and future food interaction labels. We construct two different target label types over four time horizons, for a total of eight tasks.

We denote the ground-truth food-interaction label at time *t* as ℒ (*t*). Given a history of poses from time *t − L* to *t* and a buffer period *b*, the label types are:

1. Single Frame: Forecasting the food interaction label at *t* + *b*

Definition: The label at time *t* + *b* is positive (1) if the animal is engaging with the food at time *t* + *b*, otherwise it is negative (0).

2. Transition: Forecasting a transition ℒ (*t*) = 0 *→* ℒ (*t* + *b*) = 1

Definition: Given that the animal is not engaging with the food at time *t*, the label is positive if the animal is engaging with the food at time *t* + *b*, otherwise it is negative.

The automated labeling procedure is visualized in Figure 1. We evaluate forecastability across three time horizons: 100-milliseconds, 1-second, 2-seconds, and 5-seconds. Evaluating each pair of (label-type, time-horizon) yields 8 separate binary classification tasks.

**Figure 1.**
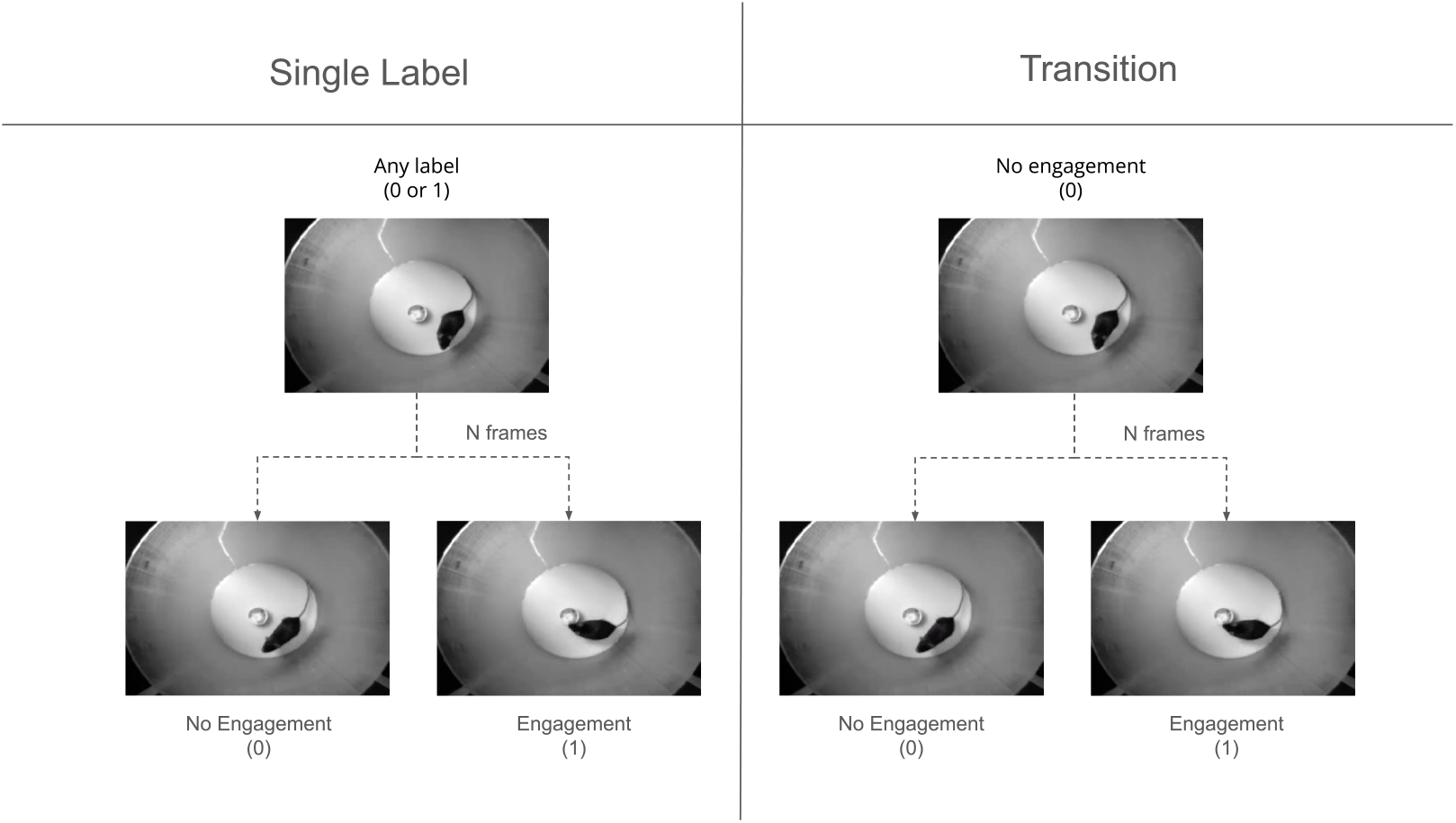
Diagram of food interaction labeling procedure. Left: single-frame target prediction. A sequence of frames is classified as positive (1) if the animal is engaging with the food after a buffer of *N* frames, otherewise it is classified as negative (0). Right: transition target prediction. A sequence of frames is classified as positive (1) if the animal is not engaging with the food at frame *t* and engaging with the food at frame *t* + *N* ; a sequence of frames is classified as negative (0) if the animal is not engaging with the food at frame *t* and also not engaging with the food at frame *t* + *N* . Any sequence of frames in which the animal is engaging with the food at frame *t* is excluded from the training/validation sets.

We also trained a model to forecast future *trajectories* of animal poses, rather than forecasting the label itself. Formally, this is a multioutput regression task over 2*n* variables: *x* and *y* coordinates for each of *n* body parts. One can convert predictions of future poses – with a buffer length *b* – into a predicted label similar to the previously-described task by determining whether the predicted snout coordinates are contained within the bowl’s radius, and evaluate the quality of the predictions in the same way as described previously. Instead of making a binary prediction, we ordered the likelihood of food interaction by predicted (normalized) Euclidean distance from the food bowl: given normalized food bowl center coordinates (*h*_*c*_, *w*_*c*_) and normalized predicted snout coordinates 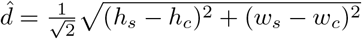 be the Euclidean distance between predicted snout coordinates and the food bowl center coordinates, normalized by a constant 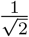 to constrain values to the unit interval [0, 1]. Then the probability of food interaction is defined as

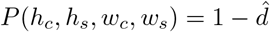

This means that we assign a probability of food interaction inversely proportional to the predicted distance. The upside of this “soft prediction” (i.e. probabilistic rather than binary) is that it allows for varying degrees of confidence, and varying-threshold classification metrics such as average precision can be used to evaluate the model’s capability. We chose the PyTorch-Forecasting implementation of the Temporal Fusion Transformer (TFT) [49, 50] as our model of choice for pose trajectory prediction. The training procedure for the TFT is described in section 2.4.2, and the overall data and training pipeline is visualized in Figure 2.

**Figure 2.**
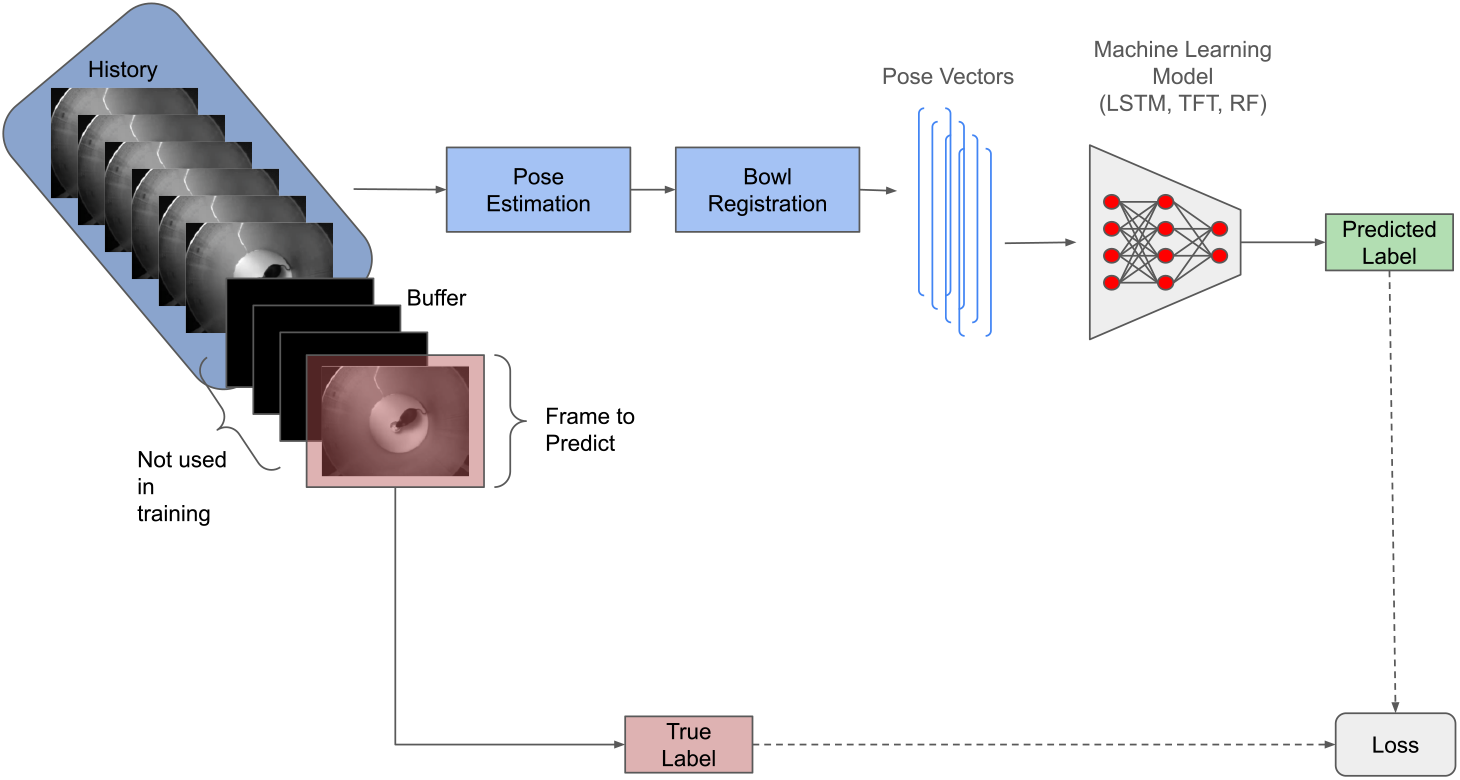
Overview of the data preprocessing and training/inference procedures. A sequence of frames is constructed with three components: (1) a history of frames, which is fed into the network, a buffer of size *N*, which consists of *N −* 1 frames over which predictions are not made, and a frame at the end of the buffer at which we predict whether or not the animal is engaged with the food. For a given sequence of video frames, we predict body-part locations for each frame with a pose estimation network, then transform the predicted coordinates to a new coordinate system whose origin is the center of the food bowl. These re-centered history coordinates are fed into a machine learning model which is trained to make a prediction on the frame at the end of the buffer. We optimize the LSTM network with a binary cross-entropy loss function between the predicted and ground-truth labels; we optimize the Random Forest with a Gini criterion; and we optimize the TFT with a mean absolute error (MAE) loss between the predicted location of the mouse’s snout coordinates and its actual coordinates at the frame to predict.

### 2.3 Random Forest Baseline

A major advantage of deep neural networks is that they can be jointly optimized with other deep neural networks. This way, if you have paired heterogeneous data types (e.g. synchronized behavioral video and calcium imaging of a neuronal population), you can train neural networks on different domains and make predictions with their combined representations of the inputs. As such, a main point of the described approach is to provide deep neural networks to learn locomotion features that encode future behaviors, with the eventual goal of using such a program in a multimodal learning paradigm incorporating other behaviorally-relevant forms of data. To assess the capability of this deep learning approach relative to other classes of more traditional machine learning approaches, we compare the LSTM and TFT performances to that of a random forest (RF), a non-differentiable ensemble tree learning algorithm that is a state-of-the-art method in statistical learning.

### 2.4 Training

#### 2.4.1 LSTM Forecasting Model

The networks are each trained using the Adam optimizer with a learning rate of 10^*−*4^, weight decay of 10^*−*4^, batch size of 256, and weighted cross-entropy loss objective (we used 0.06 : 1 negative to positive ratio due to class imbalance). For each task, we tuned hyperparameters by performing a grid search over the history length (3 seconds, 5 seconds, 10 seconds, 30 seconds), LSTM hidden vector size (32, 64, 128), and number of layers (3,4,5). Each model was trained for 500 epochs on a single NVIDIA V100 GPU.

In our experiments, we used two 30-minute videos for training and evaluation. The entirety of the first video was used for training. We split the second video into three sets: the first 0-40%, middle 40-50%, and final 50-100%. The first 40% was used for training, the final 50% was used for validation, and the remainder in between the two sets was thrown away to avoid overlap between training and validation sets. As long as the end of the training set and beginning of the validation set are separated in time by at least the history length, there is no overlap between the training and validation sets. Because rodent behavior can be highly individualized and varies from one experiment trial (i.e. video) to another, we used data from the beginning of a trial as part of the training set in order to learn trial-specific behavioral patterns.

#### 2.4.2 Temporal Fusion Transformer Trajectory Prediction

We utilized the same training and evaluation datasets for future trajectory prediction, however, instead of treating a future *event* as the target, we treat the future sequence of pose vectors as targets and train a temporal fusion transformer to forecast these future pose trajectories. For all trajectory prediction experiments, we use a 4-layer transformer model with 8 attention heads, a hidden size of 32, and 10% dropout. The model was trained for 500 epochs with the Adam optimizer at a constant learning rate of 2 *×* 10^*−*4^ and Mean Absolute Error (MAE) loss, and a batch size of 256. The MAE loss is calculated between the predicted and actual locations of the mouse body parts, with equal weight given to each body part.

#### 2.4.3 Random Forest Baseline

We use an ensemble of 500 trees, each with a maximum depth of 15. Each tree is trained on at most 5% of the samples and at most 25% of the features (features are locations of a single body-part at a single timestamp).

## 3 Results

In this section we evaluate the predictability of food interaction at multiple time-scales from behavioral video data alone. For each of the 8 prediction tasks, we train an LSTM network, along with a random forest model as a baseline. Given that there is a large class imbalance – especially for transition tasks – we report average precision score as a measure of classifier performance, which represents a tradeoff between precision and recall at varying confidence thresholds for classifying a sample as positive or negative.

Shown in Table 1 are the classification metrics for each task. We evaluated all trained LSTM network and Random Forest models across each set of hyperparameters, and present the best test-set results and hyperparameters for each task. As expected, the classifiers’ abilities degrade as the window over which to forecast gets larger. Transitions are considerably more difficult to forecast, likely because the ratio of number of transitions to number of non-transitions is much lower than the ratio of number of frames with food interactions to number of frames with non-food-interactions, presenting a challenging imbalanced learning problem for the machine learning models and generally less positive samples in the transition set to learn from.

**Table 1:** Comparison of machine learning models across different forecasting tasks. We report the Average Precision score for each type of label (single-label or transition between food-interaction states) and buffer period (100ms, 1 second, 2 seconds, 5 seconds).

| Label Type | Buffer<br>(frames \| seconds) | Average<br>Precision<br>LSTM<br>(%) | Average<br>Precision<br>Random<br>Forest<br>(%) | Average<br>Precision<br>TFT<br>(%) |
| --- | --- | --- | --- | --- |
| Single Frame | 3 \| 0.1 | 83.64 | 90.68 | **97.70** |
| Transition | 3 \| 0.1 | 8.85 | 19.41 | **54.16** |
| Single Frame | 30 \| 1 | 77.45 | 40.03 | **78.30** |
| Transition | 30 \| 1 | 22.23 | 16.89 | **27.24** |
| Single Frame | 60 \| 2 | 60.67 | 23.65 | **63.32** |
| Transition | 60 \| 2 | **20.13** | 7.93 | 18.57 |
| Single Frame | 150 \| 5 | **52.36** | 15.75 | 44.97 |
| Transition | 150 \| 5 | **25.92** | 9.60 | 15.78 |

Over shorter timescales of 100ms and 1s, the TFT is clearly the best performing model, outperforming the LSTM and RF on both single-frame and transition tasks. However, it begins to lose its dominance at the 2s timescale, as it performs about on par with the LSTM. At the 5s timescale, it clearly falls behind the LSTM, suggesting that forecasting the exact trajectory of body-parts is exceedingly challenging at such timescales when only historical locomotion activity is available.

## 4 Discussion

In this study we demonstrate the predictability of specific, meaningful animal behavioral events from properties of the motion of key body parts alone.

Based on results gathered from initial experiments, we found that the LSTM network outperformed Random Forest on longer timescales, but underperformed on smaller timescales, especially 100 milliseconds. We hypothesized that because the ratio of positive to negative samples created by our label construction method decreases with the buffer size, with a buffer size of 3 frames there are too few positive samples to learn from, thus the network is discouraged from making predictions far away from 0, even for positive samples. To overcome this, we thought that at short timescales such as 100 milliseconds, we could train a network to reliably forecast body-part trajectories; then we could convert these trajectory predictions to labels and evaluate the model’s performance in the same way that we evaluated the label forecasting method. The TFT model that we trained did indeed perform much better than the LSTM and Random Forest on the 100ms timescale, indicating the predictability of food-interaction-specific trajectories in the near-term. However, on longer timescales the TFT’s performance fell behind the LSTM; we suspect this is due to compounding errors over the long term which leads to trajectories that diverge greatly from the actual trajectory.

Because the deep learning algorithms developed in this study are end-to-end differentiable, they can be jointly optimized with other deep learning modules with other dynamic inputs, such as calcium imaging or fiber photometry recordings, embracing the rising “differentiable biology” paradigm [51]. Their representations can also be concatenated with static biological or experimental covariates, such as gene knockouts, behavioral training, etc., to enhance the learned representations. We note that due to the small parameter count and high degree of parallelization afforded by the deep learning models, they can be used in real-time behavior sessions, both for inference and within-session training of the algorithm. The relatively small parameter count and fast inference time enables our model to be used as a component in closed-loop prediction-focused neuromodulation systems, from those enabled by optogenetics to those enabled by electrode-based stimulation, or to automate behavioral training, by administering interventions such as foot shocks. Simple rule-based strategies for automating animal behavioral training exist, but training behaviors that depend on more complicated behavioral patterns still requires experimentalists’ supervision; by learning a mapping from compressed representations of experimental data such as video and physiological recordings to behaviors of interest, the paradigm introduced in this paper presents a path to automate such complicated training procedures.

This study sets the stage for further types of behavior prediction from behavioral data. Some avenues of future work include scaling up training to more animals to learn trial-agnostic behavioral features, predicting animal pose trajectories, forecasting future aggregate information on long timescales (as opposed to individual behavioral events on short timescales), and incorporating other recording modalities to learn better representations. Self-supervised pretraining with right-masked generative modeling objectives on pose sequences presents an interesting path to foundation models for animal behavior.

## 5 Acknowledgements and Funding

This work was supported in part through the computational and data resources and staff expertise provided by Scientific Computing and Data at the Icahn School of Medicine at Mount Sinai and supported by the Clinical and Translational Science Awards (CTSA) grant UL1TR004419 from the National Center for Advancing Translational Sciences.

PJK is grateful for support from the National Institute on Drug Abuse (NIDA) (grants DA053629 and DA047233) and the Cure Addiction Now (CAN) Foundation.

